# On the Accuracy of Internal Circadian Time Prediction Methods from a Single Sample

**DOI:** 10.64898/2026.02.11.705208

**Authors:** Michael T. Gorczyca

**Affiliations:** MTG Research Consulting

**Keywords:** Circadian rhythm, Dim-light melatonin onset, Machine learning, Measurement error, Transcriptome

## Abstract

Biological processes ranging from gene expression to sleep-wake cycles display oscillations with an approximately 24-hour period, or circadian rhythms. A challenge in analyzing circadian rhythms is that these rhythms vary across individuals and are based on an individual’s internal circadian time (ICT), which is uniquely offset relative to the 24-hour day-night cycle time (zeitgeber time, or ZT). Many model-based methods have been proposed to predict ICT given an individual’s biomarker measurements. However, the prediction accuracy of these methods is rarely validated using known ICT. In this article, we evaluate this accuracy for three state-of-the-art model-based methods: COFE, partial least squares regression, and TimeSignature. We find that if a single sample is obtained from each individual and a model is fit using only biomarker measurements as predictors, then ZT predicts ICT more accurately than any of the model-based ICT predictions. However, we also find that TimeSignature can outperform ZT when the model incorporates sine and cosine transforms of sample collection ZT as two additional predictors. These findings are based on analysis of three circadian transcriptome datasets as well as simulation studies, and highlight the importance of accounting for individual-level differences in biomarker oscillations to improve ICT prediction.

## 1 Introduction

Biological processes ranging from gene expression to sleep-wake cycles display oscillations with an approximately 24-hour period, or circadian rhythms (Czeisler and Gooley, 2007; Zhang et al., 2014). These rhythms are generated by the circadian clock, an internal timing system with neural and molecular mechanisms that support synchronizing the timing of these biological processes to the 24-hour day-night cycle (Bae et al., 2001; Hastings et al., 2018; Herzog et al., 2017; Yamaguchi et al., 2003). Notably, an individual’s circadian clock time (internal circadian time, or ICT) is not necessarily equal to the 24-hour day-night cycle time (zeitgeber time, or ZT). Rather, an individual’s ICT can be uniquely offset relative to ZT due to that individual’s genetic makeup (Hsu et al., 2015), age (Kennaway, 2023), and interactions with the environment (Allebrandt et al., 2014; Stothard et al., 2017; Wright et al., 2013).

Over the past two decades, increasing evidence has indicated that knowledge of an individual’s ICT has implications in healthcare (Hughes et al., 2017), from optimizing the timing of therapeutic interventions (Dallmann et al., 2014, 2016) to diagnosing circadian rhythm disorders (Zee et al., 2013). However, there has been limited adoption of these translational research discoveries into clinical practice (Dallmann et al., 2016; Nelson et al., 2024; Ruben et al., 2019). One reason for this lack of adoption is that determination of ICT in research settings can be logistically challenging. Specifically, the gold-standard measurement of the offset for an individual’s ICT relative to ZT is that individual’s dim-light melatonin onset (DLMO) time, which is defined as the ZT at which an individual begins to secrete melatonin under dim-light conditions (Lewy, 1999; Kantermann et al., 2015; Kennaway, 2019, 2020; Reid, 2019). DLMO time determination traditionally requires the collection of blood or saliva samples over an extended period of time, the availability of technicians to collect these samples, access to facilities to maintain controlled conditions, and the use of high-quality assays to obtain accurate melatonin measurements from each sample (Wittenbrink et al., 2018; Lewy, 1999; Kennaway, 2019, 2020, 2023; Hughey, 2017; Ruiz et al., 2020). These conditions are not always feasible for an investigator when performing an experiment, and compromises on DLMO time determination reduce the reliability of translational research discoveries (Gorczyca, 2026; Gorczyca et al., 2024a,b; Sollberger, 1962; Weaver and Branden, 1995).

To circumvent these challenges and support reliable discoveries, circadian biology research has increasingly focused on developing model-based methods for predicting the ICT at which a distinct set of study-relevant biomarkers are measured (Anafi et al., 2017; Ananthasubramaniam and Venkataramanan, 2025; Ogholbake and Cheng, 2025; Braun et al., 2018; Han et al., 2025; Huang and Braun, 2024; Hughey, 2017; Laing et al., 2017; Liang et al., 2020). These methods can be divided into two groups: supervised machine learning methods, which develop models that explicitly predict measurement time (either ICT or ZT) given biomarker measurements (Braun et al., 2018; Huang and Braun, 2024; Hughey, 2017; Laing et al., 2017); and unsupervised machine learning methods, which develop models that recover ICT without having any data concerning ICT or ZT (Anafi et al., 2017; Ogholbake and Cheng, 2025; Han et al., 2025; Liang et al., 2020). It is emphasized that these methods aim to predict ICT given biomarkers measured at a single time point, in part to minimize the cost of ICT determination and burden on an individual.

This article is motivated by the potential clinical utility of accurately predicting an individual’s ICT and provides a critical evaluation of existing ICT prediction methods. We show that none of the methods considered account for individual-level differences in biomarker oscillations, which can lead to ZT predicting ICT more accurately than any model-based ICT prediction. However, we also demonstrate that model-based ICT prediction accuracy generally improves when model development incorporates both biomarker measurements and the ZT of sample collection.

We organize the remainder of this article as follows. Section 2 gives an overview of our real and simulated data as well as the ICT prediction methods considered. Section 3 presents the results from applying each method on real and simulated data. Finally, Section 4 discusses the results of this study and directions for future work.

## 2 Materials and Methods

### 2.1 Data Overview

#### 2.1.1 Preliminaries and Notation for Data

For each dataset considered in this article, we assume that the data was obtained from a circadian biology experiment. Each sample was collected from a different individual, unless stated otherwise. For the *i*th individual, measurements are obtained for *G* distinct biomarkers. We denote 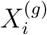 as the recorded measurement of the *g*th biomarker for individual *i*.

Each sample is associated with both a zeitgeber time (ZT), denoted as 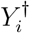 and an internal circadian time (ICT), denoted as 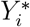.To relate ICT and ZT, we define *Z*_*i*_ as the dim-light melatonin onset (DLMO) time for individual *i*. ICT is then expressed as

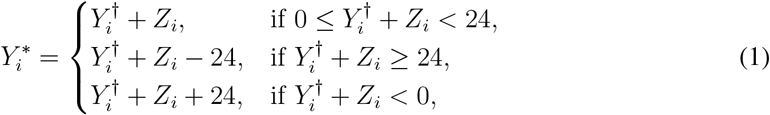

where all times are measured in hours on a 24-hour scale.

We assume that biomarker oscillations follow an individual-level multi-component cosinor model with respect to ICT (Bingham et al., 1982; Cornelissen, 2014; Tong, 1976). Specifically, the oscillations of the *g*th biomarker for individual *i* can be modeled as

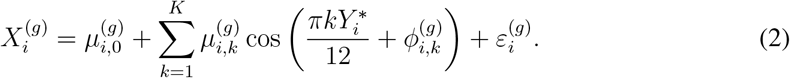

In Equation 2, 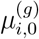 denotes the midline estimating statistic of rhythm (MESOR) for the *g*th biomarker of individual *i*. Each term in the summation corresponds to the *k*th harmonic of 24-hour oscillation. Specifically, for the *k*th harmonic, 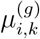 represents the oscillation amplitude, defined as the deviation from the MESOR to the harmonic peak, and 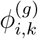 denotes the corresponding phase-shift parameter, which determines the timing of peaks for that oscillation harmonic relative to the 24-hour oscillation period (Bingham et al., 1982; Cornelissen, 2014; Gorczyca et al., 2025). The error term 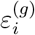 represents random measurement noise associated with the *g*th biomarker, and is assumed to have a mean of zero; no additional assumptions about the distribution of this noise are imposed here. We note that while there are *G* different biomarkers, we will assume that every biomarker has *K* oscillation harmonics. One reason for this modeling choice is that biomarkers with fewer oscillation harmonics can be represented within this framework by setting 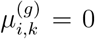 for all *k* beyond that biomarker’s oscillatory complexity. For example, if we set *K* = 3, then we would set 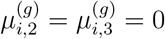 for every individual when a biomarker has one (*K* = 1) oscillation harmonic.

We emphasize that the MESOR, amplitudes, and phase-shifts of the multi-component cosinor model in Equation 2 are not assumed to be shared across individuals. Instead, all oscillation parameters are indexed by individual (*i*) and biomarker (*g*). This individual-level parameterization reflects known sources of biological and experimental variability that could reduce ICT prediction accuracy. One example to motivate this variability concerns oscillations in cortisol levels, which are known to be associated with DLMO time (Lewy and Sack, 1989; Pandi-Perumal et al., 2007) but are also influenced by meal timing, where meal timing can vary across individuals (Debono et al., 2009; Legler et al., 1982; Stimson et al., 2014). A second example is that the amplitudes of oscillation for many biomarkers decrease with age, introducing individual-level differences in these amplitudes (Hood and Amir, 2017; Nakamura et al., 2015; Wang et al., 2025). Finally, a third example from post-mortem circadian transcriptome studies involves variability in tissue sample integrity and individual-level differences in the post-mortem interval, or the time elapsed between death and post-mortem tissue sample collection (Larriba et al., 2023; Zhu et al., 2017). These individual-level differences could be interpreted as individual-level perturbations on the MESOR, amplitudes, and phase-shifts of oscillation.

#### 2.1.2 Circadian Transcriptome Data Overview and Processing

We consider data from three longitudinal transcriptome studies that are commonly used to benchmark the accuracy of ICT prediction methods (Braun et al., 2018; Huang and Braun, 2024; Hughey, 2017; Laing et al., 2017; Möller-Levet et al., 2025). In each study, multiple individuals underwent at least one experimental condition, during which two sets of blood samples were collected from each individual over time: one set was used to measure biomarkers (defined in these studies as gene expression levels), and the other was used to determine each individual’s DLMO time. We note that while these three datasets were derived from studies in which each individual contributed multiple samples over time, we will assume every sample from these three datasets is independent. This assumption has been made for ICT prediction method evaluation (Braun et al., 2018; Huang and Braun, 2024; Hughey, 2017; Laing et al., 2017) as well for illustration of circadian rhythm modeling frameworks (Gorczyca et al., 2024a; Gorczyca and Sefas, 2025).

The first dataset will be referred to as the “Archer dataset.” This dataset was generated in a study where 22 individuals participated in a four-day sleep-desynchrony protocol. Before the protocol began, blood samples were collected once every four hours over a 24-hour period to obtain pre-protocol DLMO time and biomarker data. After completing the four-day protocol, during which each individual remained awake for 20 hours and slept for eight hours in repeated cycles, blood samples were again collected from each individual once every four hours over a 24-hour period to obtain post-protocol DLMO times and biomarker data (Archer et al., 2014).

The second dataset will be referred to as the “Braun dataset.” This dataset was developed to provide an additional benchmark for evaluating the performance of different ICT prediction methods. Specifically, this dataset was created from blood samples gathered from 11 individuals with similar sleep schedules, health statuses, and ages. Each individual provided a blood sample once every two hours over a 29-hour period to obtain DLMO times and biomarker data (there is only one experimental condition) (Braun et al., 2018).

The third dataset will be referred to as the “Möller-Levet dataset.” This dataset was curated during a study on the effects of how insufficient sleep affects biomarker oscillations. The study involved 24 individuals being placed under two interventions. The first intervention allowed each individual to sleep for up to ten hours each night. The second intervention allowed each individual to sleep for up to six hours each night. After an intervention, each individual contributed a blood sample once every three hours over a 30-hour period (Möller-Levet et al., 2013).

All three datasets have been processed and made publicly available, with each processed dataset initially containing measurements from 7615 biomarkers (Huang and Braun, 2024). In this article, we perform two additional processing steps that follow the protocol of other studies (Gorczyca et al., 2024a; Gorczyca and Sefas, 2025; Gorczyca, 2026). First, data from individuals whose DLMO times could not be determined are excluded. Second, any biomarkers with missing measurements in a given dataset are removed from that dataset. A summary of each dataset after these processing steps is provided in Table 1.

**Table 1:** Summary of each circadian transcriptome dataset after data processing.

|  | **Archer** | **Braun** | **Möller-Levet** |
| --- | --- | --- | --- |
| **Total Sample Size** | 258 | 153 | 355 |
| **Number of Biomarkers** | 3689 | 7615 | 7615 |

#### 2.1.3 Simulation Data

To evaluate how individual-level differences in biomarker oscillations affects the performance of ICT prediction methods, we generate simulation data under a set of conditions based on the multicomponent cosinor model in Equation 2. In total, we consider 16 simulation settings, defined by the combination of four configuration parameters that control the complexity of biomarker oscillations and presence of individual-level differences in biomarker oscillations. There are two possible specifications for each configuration parameter, and each simulated dataset consists of *G* = 1000 biomarkers. Additional analyses that motivate our construction of these simulation configuration parameters are provided in Appendix A.

The first configuration parameter is the number of oscillation harmonics, or *K* from Equation 2. We consider *K* = 1, which corresponds to sinusoidal oscillations, and *K* = 3, which allows for more complex oscillations. In practice, circadian rhythm studies rarely specify more than three harmonics, or *K >* 3, when modeling biomarkers (Albert and Hunsberger, 2005; Hughes et al., 2009).

The second configuration parameter controls individual-level differences in oscillation amplitudes 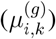 for the model in Equation 2. In one specification, amplitudes are identical across individuals, with 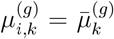 for all *i*, where 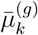 denotes a population-level amplitude for the *k*th harmonic of the *g*th biomarker. In the other specification, amplitudes vary across individuals and are generated from

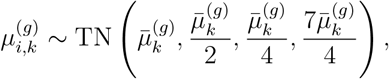

where TN(*a, b, c, d*) denotes a truncated normal distribution with mean *a*, standard deviation *b*, and bounds [*c, d*].

The third configuration parameter concerns individual-level differences in the MESOR 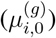 for the model in Equation 2. In one setting, all individuals share a common MESOR value, 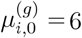 In the alternative setting, the MESOR varies across individuals according to

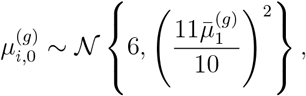

where 𝒩 (*a, b*^2^) denotes a normal distribution with a mean of *a* and a variance of *b*^2^. This distribution results in individual-level MESOR differences to be on a scale comparable to the oscillation amplitude of the first harmonic 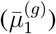 .

The fourth configuration parameter involves individual-level differences in phase-shift parameters 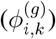 for the model in Equation 2. In one specification, phase-shifts are shared across individuals, with 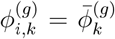 for all *i*, where 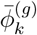 denotes a mean phase-shift for the *k*th harmonic of the *g*th biomarker across individuals. This specification implies that individual-level differences in biomarker peaks and troughs are solely due to differences in DLMO time. In the alternative specification, phase-shifts vary across individuals according to

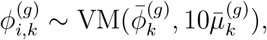

where VM(*a, b*) denotes a von Mises distribution with mean *a* and concentration *b*. The purpose of this second specification is to introduce variability in peak and trough timing that cannot be explained by ICT alone.

Each of the 16 simulation settings corresponds to a unique combination of these four configuration parameters. For each setting, we generate 500 independent simulation trials. In each trial, a dataset of *N* = 200 independent samples is generated, where the *i*th ZT is defined as

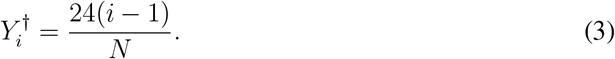

Equation 3 reflects an evenly spaced sampling design, which follows recommended protocols for circadian biology experiments (Hughes et al., 2017; Zong et al., 2023) and is known to be the optimal sample collection protocol for the multi-component cosinor model in Equation 2 based on multiple statistical criteria (Federov, 1972; Gorczyca and Sefas, 2025; Pukelsheim, 2006).

Across all 16 simulation settings, we generate the remaining model parameters as follows. Population-level amplitudes are generated as

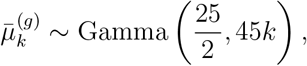

where Gamma(*a, b*) denotes a gamma distribution with shape *a* and rate *b*. Population-level phaseshifts are generated as

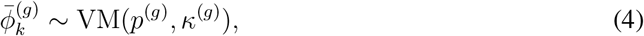

where there is a 50% chance that *p*^(*g*)^ =*−π/*2 and *κ*^(*g*)^ = 7, and a 50% chance that *p*^(*g*)^ = *π/*2 and *κ*^(*g*)^ = 14. DLMO times are generated as

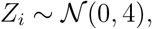

which is due to a standard deviation of two hours commonly being reported for DLMO time across cohorts (Kennaway, 2023). Finally, random measurement noise is generated as

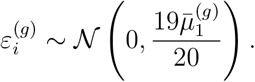

Details motivating these distributional choices for 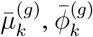 and 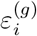 are also provided in Appendix A.

### 2.2 Prediction Method Overview

ICT prediction methods generally assume that one can map biomarker measurements to a transformation of ICT, where this transformation of ICT is defined as the two-dimensional vector

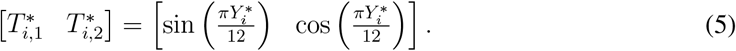

The reason that these methods consider predicting this transformation of ICT in Equation 5 is to account for nonlinearities in 24-hour time. For example, while 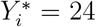 hours and 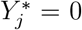 hours represent the same time point in a 24-hour period, their arithmetic difference is 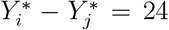 hours. The transformation in Equation 5 accounts for this behavior and enables application of traditional model development frameworks. It is also noted that ICT can be recovered from 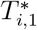 and 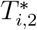 by computing

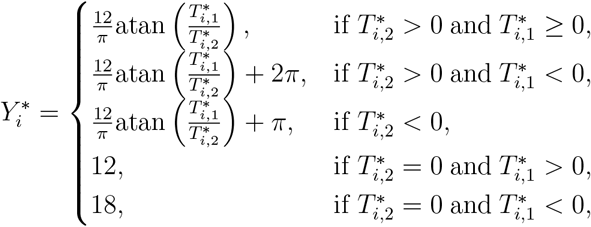

where atan(*a*) is the arctangent function with argument *a*.

When mapping biomarker measurements to transformed ICT, an investigator will assume that this mapping is based on a function, 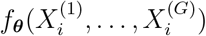 which is parameterized by a set of weights ***θ***. This function satisfies the relationship

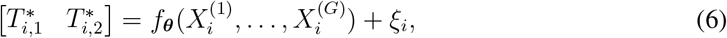

where *ξ*_*i*_ denotes random noise associated with the outputs of this function. When the function 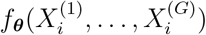 is fit using supervised machine learning, the set of weights ***θ*** would be estimated by minimizing a cost function. This estimation process would be formulated as

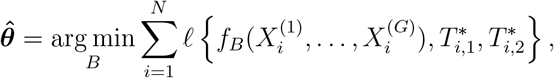

Where 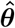 denotes the set of estimated weights, and 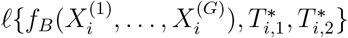 denotes the cost function relative to the transformed ICT predictions output by 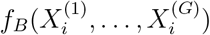 and true transformed ICT. For example, this cost function can be defined as minimizing the sum of squared errors between transformed ICT predictions and the corresponding true transformed ICT. In this article, we consider two state-of-the-art supervised machine learning methods for ICT prediction that follow this framework: partial least squares regression (Laing et al., 2017) and TimeSignature (Braun et al., 2018). These two methods have consistently obtained strong predictive performance and outperformed other supervised machine learning methods (Braun et al., 2018; Huang and Braun, 2024).

For unsupervised machine learning, ICT is instead considered an unobserved latent variable that is unavailable for model development. As a consequence, ***θ*** cannot be estimated by minimizing a cost function with respect to known ICT. Instead, unsupervised machine learning methods operate under the assumption that there is additional structure in the data that can enable recovery of ICT. In general, ICT prediction with unsupervised machine learning methods can be characterized by jointly estimating the weights of the function 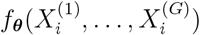 and the weights of a function 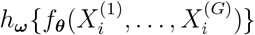 or computing

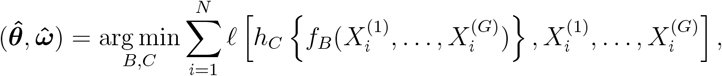

where 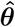 and 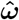denote the estimated sets of weights for ***θ*** and ***ω***, respectively. In this context, 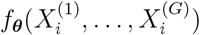 serves as a mapping of biomarker measurements to latent ICT, and 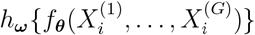 aims to reconstruct the biomarker measurements from the output of 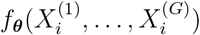 (Anafi et al., 2017; Ananthasubramaniam and Venkataramanan, 2025; Liang et al., 2020). In this article, we only consider the unsupervised machine learning method COFE, a nonlinear principal component analysis method, as COFE has outperformed other unsupervised machine learning methods (Ananthasubramaniam and Venkataramanan, 2025).

### 2.3 Model Development and Assessment

For all analyses, whether using the circadian transcriptome datasets described in Section 2.1.2 or the simulated data described in Section 2.1.3, model development and assessment follow a structured protocol. Specifically, each dataset is randomly divided into two non-overlapping groups, a training dataset and a test dataset, with 50% of the samples allocated to each. The training dataset is used to fit a model and the test datset is used to provide an unbiased assessment of predictive performance.

Each ICT prediction method (COFE, partial least squares regression, and TimeSignature) depends on one or more hyper-parameters, which are user-specified quantities that influence how a machine learning model estimates relationships between biomarker measurements and transformed ICT (Bergstra and Bengio, 2012). For each method, we consider the same sets of hyper-parameters as defined in each method’s corresponding study. To identify the optimal hyper-parameters from all hyper-parameters considered, we perform ten-fold cross-validation on the training dataset. In ten-fold cross-validation, the training dataset is divided into ten approximately equal subsets of non-overlapping samples (folds). Nine folds are used to fit a model, and the remaining tenth fold is used to evaluate predictive performance (this tenth fold can be considered a test dataset within the training dataset, which is referred to as a validation dataset). This process is repeated ten times, with each fold used once as the validation dataset, and the optimal hyper-parameters are defined as the ones that maximize a predictive performance measure of interest across all ten validation datasets (Hastie et al., 2009, Section 7.10).

Once optimal hyper-parameters are determined, each model is re-fit using the entire training dataset with these optimal hyper-parameters specified, and predictive performance is assessed on the test dataset. For the real data analyses, this procedure is repeated 500 times (500 trials are performed), each time with a randomly defined training dataset and test dataset to account for variability in how the entire dataset could be partitioned. For the simulation studies, this procedure was performed once each simulation trial, as each simulation trial generates a new dataset.

Predictive performance is quantified using three performance measures for partial least squares regression and TimeSignature. The first performance measure is the mean squared error (MSE), which is the average squared difference between predicted and true ICT. Specifically,

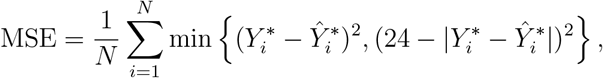

Where 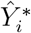 denotes an ICT prediction, which is computed with the identity in Equation 1 using transformed ICT prediction outputs from a model. For the MSE, smaller values indicate better predictive performance. We note that the error term is defined as 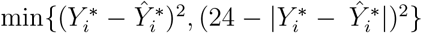 to obtain the smallest error between predicted and true ICT. For example, the difference between 23 hours and 1 hour would be 2 hours, not 22 hours. The second performance measure is the median absolute error (MedAE), which is defined as

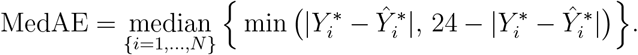

The MedAE can be considered a robust summary measure of prediction error, as a small number of samples with poor predictive accuracy do not distort its numeric value. Smaller MedAE values also indicate better predictive performance. Finally, the third measure is the absolute value of Pearson’s correlation coefficient between predicted and true ICTs, which quantifies the strength of association between these two quantities and ranges from zero to one in value. Larger numeric values indicate better predictive performance. In Appendix B, we briefly describe how we compute Pearson’s correlation coefficient with predicted and true ICT.

In this article, we select the hyper-parameters that minimized the MSE for 10-fold crossvalidation for partial least squares regression and TimeSignature, and save the ICT predictions output by these two methods on the test dataset at the end of each trial. For COFE, we instead select the hyper-parameters that maximize Pearson’s correlation coefficient for 10-fold cross-validation, and only compute this quantity for COFE on the test dataset. The reason for this decision is that COFE is based on principal component analysis (PCA). In PCA-based methods, ICT predictions are identifiable only up to an arbitrary scaling factor, which implies that assessing COFE using MSE or MedAE could be misleading (Jolliffe, 2002, Section 3.6).

We note that, for each trial, each method will undergo this model development and assessment protocol twice. The first time a method undergoes this protocol, model development will be performed using only the biomarker measurements as predictors (input variables). The second time a method undergoes this protocol, model development will be performed using the biomarker measurements as predictors together with two additional ZT-based predictors. Specifically, these two additional ZT-based predictors are defined as

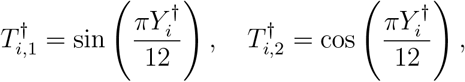

where this transformation of ZT is based on based on Equation 5 and is motivated by the fact that ICT is only offset relative to ZT in Equation 1. We also note that, to provide a baseline for assessing each ICT prediction method, we compute the MSE, MedAE, and the absolute value of Pearson’s correlation coefficient between true ICT and ZT. This baseline corresponds to using ZT as the ICT prediction, or assuming that ICT and ZT are equal across all samples. These three performance measures are computed using the test dataset in each trial, with ZT substituted for the predicted ICT.

## 3 Results

### 3.1 Real Data

Table 2 presents the results of evaluating each method on the circadian transcriptome datasets discussed in Section 2.1.2. The methods were assessed based on three performance measures: mean squared error (MSE), median absolute error (MedAE), and Pearson’s correlation coefficient (CC) on test datasets that consist of samples not used for model development. Overall, the baseline of ZT as an ICT prediction consistently outperforms all model-based ICT predictions in terms of the MSE and CC. This is not the case for MedAE, however, as partial least squares regression and TimeSignature obtain a smaller MedAE than baseline ZT on the Braun dataset. When ZT is available for model development, we find that the inclusion of additional ZT-based predictors consistently improves predictive performance for partial least squares regression and TimeSignature. In particular, TimeSignature performs at least as well as baseline ZT in terms of the MedAE on all three datasets.

**Table 2:** Predictive performance of each method on three circadian transcriptome datasets (Archer, Braun, and Möller-Levet). “PLSR” denotes partial least squares regression, and “NA” denotes not applicable. Median test dataset values across 500 simulation trials are reported, with corresponding standard deviations shown in parentheses. For each dataset and performance measure, results that perform at least as well as the ZT baseline are shown in bold.

| Archer Dataset |  |  |  |  |
| --- | --- | --- | --- | --- |
| Type | Method | MedAE | MSE | CC |
| Baseline | ZT | 1.883 (0.113) | 4.860 (0.289) | 0.971 (0.003) |
| Biomarkers Only | TimeSignature | 2.380 (0.240) | 15.570 (2.163) | 0.900 (0.016) |
|  | PLSR | 2.278 (0.260) | 16.181 (2.544) | 0.874 (0.020) |
|  | CF | NA | NA | 0.726 (0.040) |
| Biomarkers and ZT | TimeSignature | **1.645 (0.161)** | 7.944 (1.402) | 0.929 (0.012) |
|  | PLSR | 2.011 (0.258) | 12.553 (2.453) | 0.893 (0.021) |
|  | CF | NA | NA | 0.726 (0.042) |
| Braun Dataset |  |  |  |  |
| Type | Method | MedAE | MSE | CC |
| Baseline | ZT | 2.000 (0.013) | 3.930 (0.232) | 0.981 (0.002) |
| Biomarkers Only | TimeSignature | **1.711 (0.221)** | 7.717 (1.857) | 0.916 (0.019) |
|  | PLSR | **1.776 (0.233)** | 7.491 (1.923) | 0.930 (0.017) |
|  | CF | NA | NA | 0.689 (0.055) |
| Biomarkers and ZT | TimeSignature | **1.323 (0.163)** | 4.487 (1.321) | 0.949 (0.014) |
|  | PLSR | **1.711 (0.230)** | 7.342 (1.989) | 0.929 (0.018) |
|  | CF | NA | NA | 0.684 (0.053) |
| Möller-Levet Dataset |  |  |  |  |
| Type | Method | MedAE | MSE | CC |
| Baseline | ZT | 1.133 (0.046) | 2.984 (0.184) | 0.967 (0.003) |
| Biomarkers Only | TimeSignature | 1.272 (0.122) | 5.406 (0.899) | 0.943 (0.010) |
|  | PLSR | 1.228 (0.124) | 5.971 (1.130) | 0.940 (0.011) |
|  | CF | NA | NA | 0.690 (0.047) |
| Biomarkers and ZT | TimeSignature | **1.133 (0.108)** | 3.842 (0.498) | 0.957 (0.006) |
|  | PLSR | 1.177 (0.112) | 4.723 (0.873) | 0.950 (0.009) |
|  | CF | NA | NA | 0.688 (0.052) |

Figures 1, 2, and 3 present scatter plots that compare ZT, model-based ICT predictions, and true ICT for the Archer, Braun, and Möller-Levet datasets, respectively. Each point in these scatter plots represents the average ICT prediction for a sample from a given method, weighted by the number of times that sample appeared in the test dataset, relative to the corresponding true ICT. These plots show that the absolute difference between ZT and true ICT is never greater than 4 hours across any sample. In contrast, the difference between model-based ICT predictions and true ICT exceeds four hours on all three datasets. These larger deviations provide a qualitative explanation for why ZT consistently outperforms model-based ICT predictions in terms of MSE and CC.

**Figure 1:**
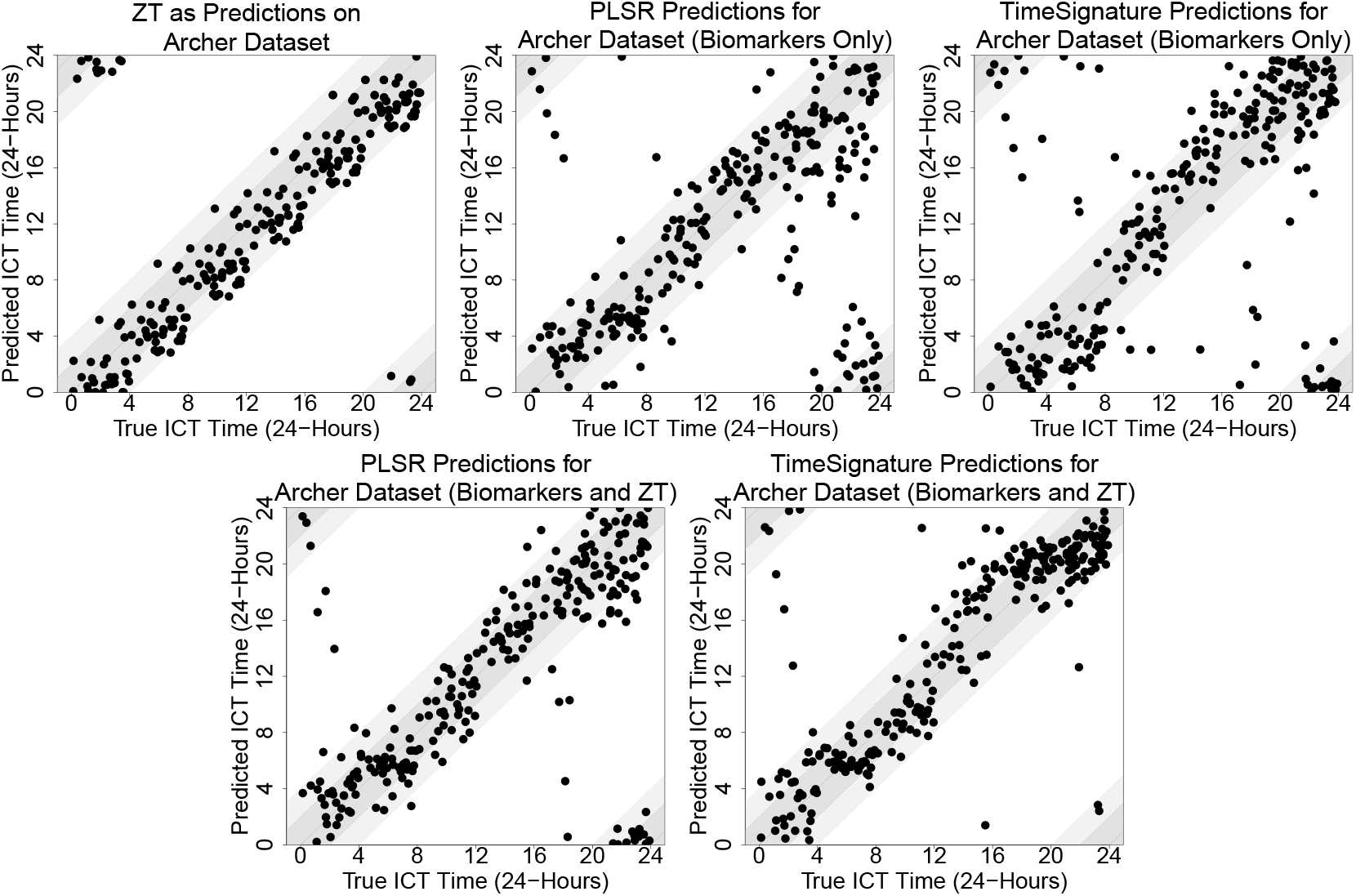
Scatter plots of internal circadian time (ICT) predictions versus true ICT for the Archer dataset produced by partial least squares regression (PLSR), TimeSignature, and baseline zeitgeber time (ZT). For each sample, the predicted ICT is obtained by computing a circular average for the predictions made when that sample appeared in the test dataset across repeated random train-test splits. The identity line represents perfect prediction, the dark gray shaded band indicates a prediction error no greater than two hours, and the light gray shaded band indicates a prediction error no greater than four hours.

**Figure 2:**
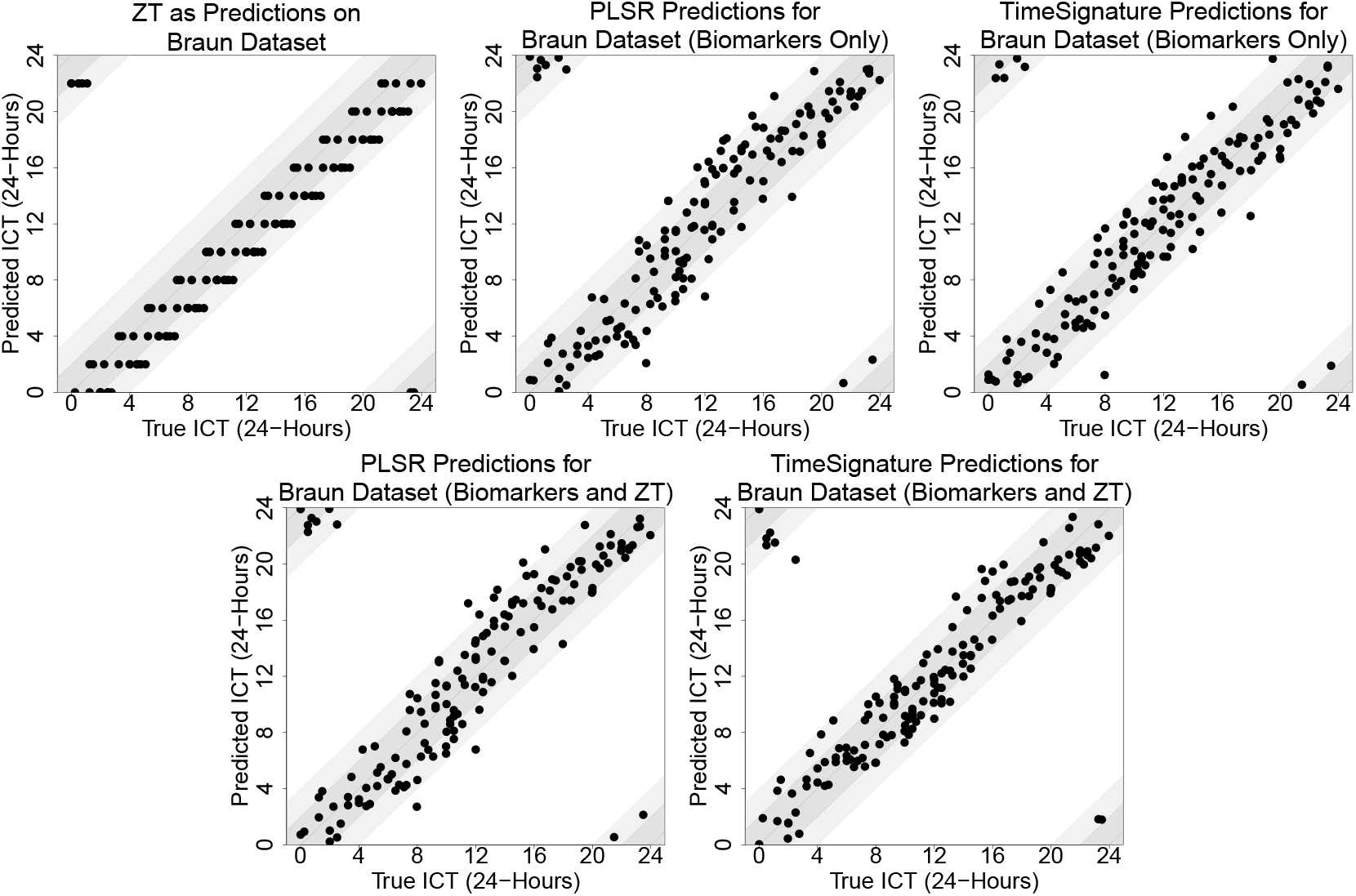
Scatter plots of internal circadian time (ICT) predictions versus true ICT for the Braun dataset produced by partial least squares regression (PLSR), TimeSignature, and baseline zeitgeber time (ZT). For each sample, the predicted ICT is obtained by computing a circular average for the predictions made when that sample appeared in the test dataset across repeated random train-test splits. The identity line represents perfect prediction, the dark gray shaded band indicates a prediction error no greater than two hours, and the light gray shaded band indicates a prediction error no greater than 4 hours.

**Figure 3:**
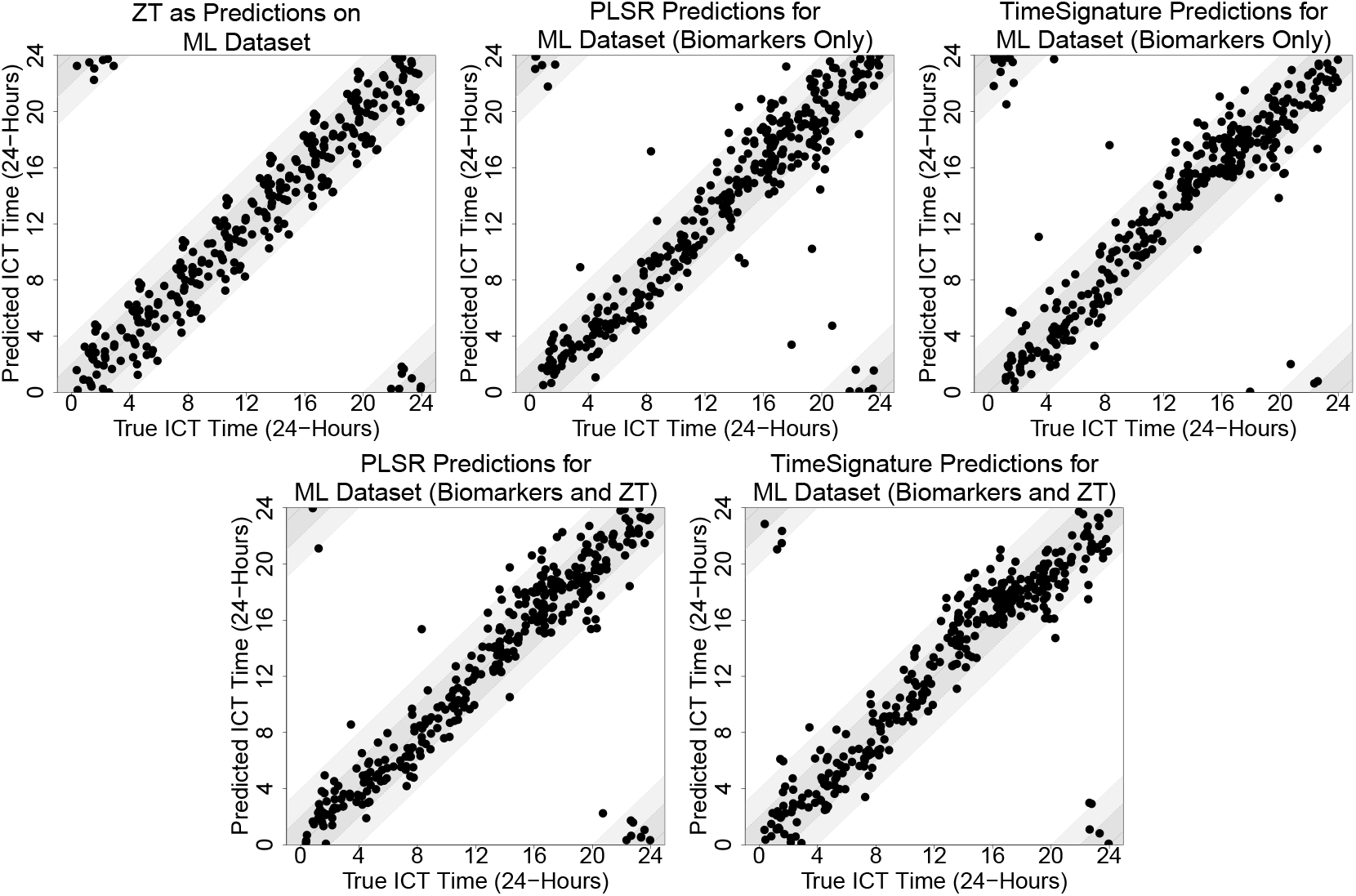
Scatter plots of internal circadian time (ICT) predictions versus true ICT for the Möller-Levet dataset produced by partial least squares regression (PLSR), TimeSignature, and baseline zeitgeber time (ZT). For each sample, the predicted ICT is obtained by computing a circular average for the predictions made when that sample appeared in the test dataset across repeated random train-test splits. The identity line represents perfect prediction, the dark gray shaded band indicates a prediction error no greater than two hours, and the light gray shaded band indicates a prediction error no greater than 4 hours.

### 3.2 Simulated Data

Table 3 summarizes ICT prediction performance results across all 16 simulation settings, where performance is again evaluated using MSE, MedAE, and CC for each method. When there are no individual-level differences in biomarker oscillation, model-based ICT prediction methods consistently outperform baseline ZT across all three performance measures. We also find that, when there are no individual-level differences, the inclusion of additional ZT-based predictors provide minimal improvement in performance. These same trends hold when only individual-level differences in the amplitudes of oscillation are present.

**Table 3:** Predictive performance of each method in each simulation setting. “‘PLSR” denotes partial least squares regression, and “NA” denotes not applicable. Median test dataset values across 500 simulation trials are reported, with corresponding standard deviations shown in parentheses.

| Simulation Setting Configurations and Performance |  |  |  |  | Baseline | Biomarkers Only |  |  | Biomarkers and Transformed Zeitgeber Time |  |  |
| --- | --- | --- | --- | --- | --- | --- | --- | --- | --- | --- | --- |
| $K$ | Same Amps? | Same MESOR? | Same PS? | Performance Measure | Zeitgeber Time | COFE | PLSR | TimeSignature | COFE | PLSR | TimeSignature |
| $K = 1$ | Yes | Yes | Yes | MSE | 1.986 (0.272) | NA | 0.701 (0.197) | 0.945 (0.258) | NA | 0.556 (0.140) | 0.566 (0.110) |
|  |  |  |  | MedAE | 0.964 (0.110) | NA | 0.481 (0.073) | 0.562 (0.081) | NA | 0.443 (0.068) | 0.462 (0.057) |
|  |  |  |  | CC | 0.980 (0.003) | 0.768 (0.081) | 0.993 (0.002) | 0.990 (0.003) | 0.766 (0.084) | 0.994 (0.001) | 0.994 (0.001) |
|  |  |  | No | MSE | 1.990 (0.274) | NA | 1.058 (0.231) | 1.665 (0.360) | NA | 0.839 (0.182) | 1.003 (0.167) |
|  |  |  |  | MedAE | 0.960 (0.108) | NA | 0.673 (0.086) | 0.837 (0.111) | NA | 0.615 (0.084) | 0.670 (0.080) |
|  |  |  |  | CC | 0.980 (0.003) | 0.771 (0.073) | 0.990 (0.002) | 0.983 (0.004) | 0.778 (0.075) | 0.992 (0.002) | 0.990 (0.002) |
|  |  | No | Yes | MSE | 1.961 (0.293) | NA | 2.145 (0.775) | 3.119 (0.987) | NA | 1.270 (0.382) | 1.007 (0.175) |
|  |  |  |  | MedAE | 0.951 (0.112) | NA | 0.771 (0.115) | 0.924 (0.128) | NA | 0.647 (0.096) | 0.640 (0.076) |
|  |  |  |  | CC | 0.980 (0.003) | 0.763 (0.073) | 0.978 (0.008) | 0.968 (0.010) | 0.765 (0.073) | 0.987 (0.004) | 0.990 (0.002) |
|  |  |  | No | MSE | 1.978 (0.276) | NA | 3.026 (0.860) | 4.655 (1.208) | NA | 1.901 (0.489) | 1.480 (0.234) |
|  |  |  |  | MedAE | 0.958 (0.106) | NA | 1.101 (0.156) | 1.306 (0.176) | NA | 0.922 (0.124) | 0.811 (0.097) |
|  |  |  |  | CC | 0.980 (0.003) | 0.767 (0.066) | 0.969 (0.009) | 0.951 (0.013) | 0.782 (0.072) | 0.981 (0.005) | 0.985 (0.003) |
|  | No | Yes | Yes | MSE | 1.976 (0.278) | NA | 0.775 (0.234) | 1.030 (0.288) | NA | 0.604 (0.150) | 0.601 (0.119) |
|  |  |  |  | MedAE | 0.946 (0.110) | NA | 0.506 (0.082) | 0.593 (0.081) | NA | 0.457 (0.076) | 0.474 (0.058) |
|  |  |  |  | CC | 0.980 (0.003) | 0.764 (0.086) | 0.992 (0.002) | 0.990 (0.003) | 0.772 (0.084) | 0.994 (0.002) | 0.994 (0.001) |
|  |  |  | No | MSE | 1.988 (0.285) | NA | 1.163 (0.244) | 1.711 (0.413) | NA | 0.922 (0.184) | 1.005 (0.180) |
|  |  |  |  | MedAE | 0.955 (0.110) | NA | 0.707 (0.094) | 0.855 (0.116) | NA | 0.636 (0.086) | 0.657 (0.081) |
|  |  |  |  | CC | 0.980 (0.003) | 0.769 (0.072) | 0.989 (0.003) | 0.983 (0.005) | 0.772 (0.075) | 0.991 (0.002) | 0.990 (0.002) |
|  |  | No | Yes | MSE | 1.978 (0.281) | NA | 2.293 (0.882) | 3.189 (1.066) | NA | 1.334 (0.432) | 1.023 (0.177) |
|  |  |  |  | MedAE | 0.958 (0.109) | NA | 0.789 (0.126) | 0.933 (0.133) | NA | 0.667 (0.099) | 0.635 (0.078) |
|  |  |  |  | CC | 0.980 (0.003) | 0.758 (0.069) | 0.976 (0.009) | 0.967 (0.011) | 0.764 (0.077) | 0.986 (0.004) | 0.990 (0.002) |
|  |  |  | No | MSE | 1.998 (0.280) | NA | 2.907 (0.802) | 4.239 (1.064) | NA | 1.893 (0.434) | 1.410 (0.216) |
|  |  |  |  | MedAE | 0.950 (0.108) | NA | 1.067 (0.149) | 1.245 (0.161) | NA | 0.901 (0.119) | 0.782 (0.090) |
|  |  |  |  | CC | 0.980 (0.003) | 0.765 (0.069) | 0.971 (0.009) | 0.956 (0.012) | 0.769 (0.067) | 0.981 (0.005) | 0.986 (0.003) |
| $K = 3$ | Yes | Yes | Yes | MSE | 1.993 (0.272) | NA | 1.782 (0.722) | 2.390 (0.870) | NA | 1.142 (0.376) | 0.920 (0.162) |
|  |  |  |  | MedAE | 0.958 (0.106) | NA | 0.719 (0.105) | 0.846 (0.115) | NA | 0.625 (0.090) | 0.600 (0.074) |
|  |  |  |  | CC | 0.980 (0.003) | 0.761 (0.077) | 0.982 (0.008) | 0.976 (0.009) | 0.767 (0.073) | 0.988 (0.004) | 0.991 (0.002) |
|  |  |  | No | MSE | 1.972 (0.263) | NA | 2.375 (0.722) | 3.509 (1.022) | NA | 1.552 (0.383) | 1.273 (0.199) |
|  |  |  |  | MedAE | 0.951 (0.106) | NA | 0.941 (0.127) | 1.121 (0.146) | NA | 0.811 (0.112) | 0.739 (0.088) |
|  |  |  |  | CC | 0.980 (0.003) | 0.772 (0.069) | 0.976 (0.008) | 0.964 (0.011) | 0.768 (0.068) | 0.984 (0.004) | 0.987 (0.002) |
|  |  | No | Yes | MSE | 1.979 (0.272) | NA | 5.107 (1.749) | 6.561 (1.948) | NA | 2.394 (0.795) | 1.346 (0.217) |
|  |  |  |  | MedAE | 0.945 (0.111) | NA | 1.101 (0.178) | 1.294 (0.183) | NA | 0.871 (0.133) | 0.759 (0.092) |
|  |  |  |  | CC | 0.980 (0.003) | 0.764 (0.068) | 0.948 (0.018) | 0.934 (0.020) | 0.756 (0.071) | 0.975 (0.008) | 0.986 (0.002) |
|  |  |  | No | MSE | 1.971 (0.274) | NA | 5.819 (1.851) | 7.737 (1.929) | NA | 2.940 (0.877) | 1.580 (0.238) |
|  |  |  |  | MedAE | 0.958 (0.113) | NA | 1.356 (0.200) | 1.580 (0.219) | NA | 1.056 (0.142) | 0.828 (0.097) |
|  |  |  |  | CC | 0.980 (0.003) | 0.766 (0.061) | 0.938 (0.020) | 0.918 (0.021) | 0.770 (0.067) | 0.970 (0.009) | 0.984 (0.003) |
|  | No | Yes | Yes | MSE | 1.967 (0.273) | NA | 1.925 (0.739) | 2.598 (0.820) | NA | 1.215 (0.394) | 0.957 (0.166) |
|  |  |  |  | MedAE | 0.946 (0.111) | NA | 0.732 (0.116) | 0.878 (0.121) | NA | 0.634 (0.102) | 0.614 (0.080) |
|  |  |  |  | CC | 0.980 (0.003) | 0.760 (0.074) | 0.980 (0.008) | 0.973 (0.009) | 0.767 (0.079) | 0.988 (0.004) | 0.990 (0.002) |
|  |  |  | No | MSE | 1.975 (0.277) | NA | 2.534 (0.800) | 3.458 (0.969) | NA | 1.591 (0.452) | 1.190 (0.189) |
|  |  |  |  | MedAE | 0.951 (0.099) | NA | 0.932 (0.129) | 1.085 (0.143) | NA | 0.789 (0.109) | 0.719 (0.085) |
|  |  |  |  | CC | 0.980 (0.003) | 0.762 (0.067) | 0.974 (0.009) | 0.965 (0.011) | 0.775 (0.070) | 0.984 (0.005) | 0.988 (0.002) |
|  |  | No | Yes | MSE | 1.986 (0.269) | NA | 5.125 (1.743) | 6.555 (1.841) | NA | 2.372 (0.750) | 1.343 (0.212) |
|  |  |  |  | MedAE | 0.956 (0.110) | NA | 1.098 (0.181) | 1.293 (0.186) | NA | 0.872 (0.134) | 0.757 (0.089) |
|  |  |  |  | CC | 0.980 (0.003) | 0.761 (0.069) | 0.947 (0.018) | 0.933 (0.019) | 0.759 (0.070) | 0.976 (0.008) | 0.986 (0.002) |
|  |  |  | No | MSE | 1.963 (0.280) | NA | 6.052 (1.752) | 7.614 (1.946) | NA | 3.030 (0.849) | 1.503 (0.235) |
|  |  |  |  | MedAE | 0.954 (0.108) | NA | 1.321 (0.193) | 1.530 (0.210) | NA | 1.044 (0.143) | 0.815 (0.096) |
|  |  |  |  | CC | 0.980 (0.003) | 0.768 (0.064) | 0.936 (0.019) | 0.921 (0.021) | 0.770 (0.067) | 0.969 (0.010) | 0.984 (0.003) |

When individual-level differences in the MESOR or phase-shifts of oscillation are introduced, predictive performance declines for all methods. Further, ZT is more accurate in ICT prediction than all other model-based ICT predictions. We note that, when additional ZT-based predictors are included for model development, the MSE, MedAE, and CC improve for all three methods. We also note that the performance measures obtained when there are individual-level differences in the MESOR and phase-shifts appear to align with the results from the real datasets in Section 3.1.

## 4 Discussion

In this article, we evaluate the performance of three state-of-the-art ICT prediction methods, COFE, partial least squares regression, and TimeSignature, using both real circadian transcriptome datasets and simulated data. We found that ignoring individual-level differences in biomarker oscillations can harm model-based ICT prediction accuracy, to the extent that ZT is more accurate in predicting ICT than any model-based ICT prediction. The simulation study results in Section 3.2 imply that the poor performance of these three methods could be attributed to individual-level differences in the MESOR or phase-shifts of oscillation.

A straightforward solution to improve predictive performance is to incorporate ZT into model development. We found that adding ZT-related information as additional predictors generally improved predictive performance on both the real and simulated datasets. In particular, when ZT-based predictors are used for model development with TimeSignature, the resulting ICT predictions obtained a smaller median absolute error (MedAE) than ZT. We attribute this improvement to TimeSignature’s underlying algorithm. Unlike COFE and partial least squares regression, which apply data transformations before model development, TimeSignature does not transform the data. This allows TimeSignature to directly leverage the signal from ZT-based predictors, which appears to be less effectively captured by the other methods.

This article offers multiple directions for future work. One direction concerns determining how to address individual-level differences in biomarker oscillations during model development. While these individual-level differences have been addressed in statistical modeling when ICT is unknown (Gorczyca, 2026; Gorczyca et al., 2024a), further work is needed to integrate these differences into ICT prediction models. A second direction involves investigating how individual-level differences in biomarker oscillations could impact statistical analysis of circadian rhythms. Recent research has highlighted that circadian transcriptome study conclusions could have low reproducibility (Brooks et al., 2023). This finding could, in part, be due to differences in biomarker oscillations across individuals. Lastly, a third direction is the extension of ICT prediction methods to scenarios where the oscillation period of biomarkers is unknown, which could complicate statistical analysis of biomarker data (Silverthorne et al., 2025).

## Data Availability Statement

The author has made the code scripts used to produce the results in Section 3.1 available at https://bitbucket.org/michaelgorczyca/single_sample_evaluation/.

## A Analysis of Real Data to Inform Simulation Study Design

### A.1 Setup

To inform the design of the simulation studies, we first analyze the three circadian transcriptome datasets described in Section 2.1.2. The goal of this analysis is to quantify the extent of individual-level variation in biomarker oscillations and to identify realistic distribution parameter values for simulating MESORs, oscillation amplitudes, phase-shifts, and random noise for biomarkers.

#### A.1.1 Individual-Level Cosinor Modeling

For each dataset, we fit an individual-level single-component cosinor model (*K* = 1 for the model in Equation 2) separately for each biomarker and each individual, as multiple samples were collected longitudinally for each individual. The parameters for every individual-level model are estimated using least squares regression, yielding individual-level estimates of the MESOR (denoted as 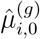oscillation amplitude 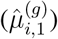 phase-shift 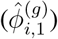 and residual variance. The residual variance is defined as

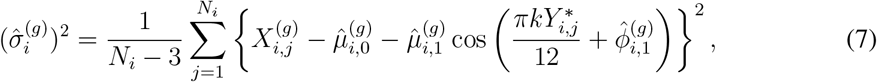

where *N*_*i*_ denotes the number of samples available for individual *i*. The denominator in Equation 7 is due to the loss of three degrees of freedom (*N*_*i*_ *−*3) when estimating the parameters of the single-component cosinor model.

#### A.1.2 Identification of Oscillating Biomarkers

To assess whether the *g*th biomarker oscillates for the *i*th individual, we assess the null hypothesis that 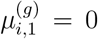 and record the resulting *p*-value. As biomarker oscillations can vary across individuals, we then use Stouffer’s method to combine the *p*-values obtained for each *g*th biomarker across every individual (Stouffer et al., 1949). After we obtain a single, combined *p*-value for each biomarker, we adjust every combined *p*-value using Bonferroni correction across all *G* biomarkers. Only biomarkers with adjusted *p*-values less than 0.05 are retained for subsequent analyses. This step ensured that downstream analyses focused on biomarkers that oscillate across individuals.

#### A.1.3 Population-Level Parameter Estimation

For each biomarker identified in Section A.1.2, we use their individual-level parameter estimates to obtain corresponding population-level quantities. To be precise, let *M* denote the number of individuals in a dataset. We define the population-level MESOR estimate 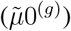 and amplitude 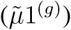 as

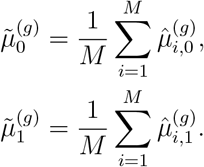

As the phase-shift is a circular quantity, the population-level phase shift is computed as the circular mean,

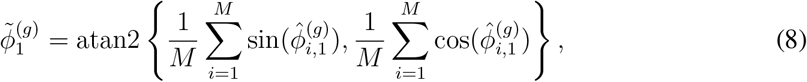

with atan2(*a, b*) denoting the two-argument arctangent function with arguments *a* and *b* (Mardia and Jupp, 1999, Equation 2.2.4). The population-level residual variance is defined as the mean of the individual-level residual variances,

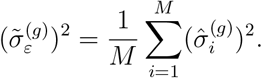

To quantify variability in the MESOR, amplitude, and phase-shift across individuals for each selected biomarker, we also computed estimates of their variance. For the MESOR and amplitude, we computed this variance as

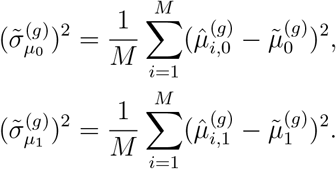

For the phase-shift, we assumed that individual-level phase-shifts followed a von Mises distribuion,

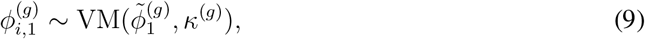

and estimated the concentration parameter *κ*^(*g*)^ using maximum likelihood methods implemented in the circular package for the R statistical software (Agostinelli and Lund, 2025). Larger values of the estimate 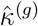 indicate stronger phase-shift concentration (less variability) across individuals.

#### A.1.4 Quantifying the Relationship between Population-Level Parameters

After we compute the quantities described in Section A.1.3 for each selected biomarker, we examine how each quantity relates to the corresponding population-level amplitude. Specifically, we performed five least squares linear regressions, each with the population-level amplitude 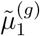 as the predictor (input variable) and one of the following quantities as the response (output variable):

1. The standard deviation of amplitude estimates, 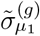
2. The standard deviation of the noise variance, 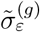
3. The range of individual-level amplitude estimates, 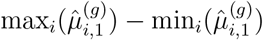
4. The standard deviation of MESOR estimates, 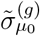
5. The von Mises concentration parameter for phase-shifts, 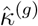

All regressions are performed without an intercept term and produced a single parameter estimate, 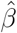. These analyses characterize how the quantities described in Section A.1.3 relate to each other, with the population-level amplitude used as a reference point.

#### A.1.5 Distribution of Population-Level Amplitudes and Phase-Shifts

Finally, to summarize how population-level amplitudes and phase-shifts are distributed, we fit distributions to each of them. For the population-level amplitude 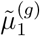, we fit a gamma distribution using maximum likelihood estimation. This distribution is used to inform specification of the population-level amplitude distribution for the simulation studies.

For the population-level phase-shifts 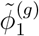we assumed they could be modeled by a two-component mixture of von Mises-Fisher distributions, as prior studies have reported that the phase-shifts qualitatively show this distribution (Korenčič et al., 2014; Rowicka et al., 2007). We compute the parameters of this two-component mixture of distributions using the movMF package in the R statistical software (Hornik and Grün, 2014) This distribution is similarly used to inform specification of the population-level phase-shift distribution for the simulation studies.

### A.2 Results

Figure 4 summarizes the relationship between population-level amplitude estimates and variability in individual-level single-component cosinor model parameters across biomarkers. Each point in this figure corresponds to a single biomarker. Application of the biomarker selection protocol described in Section A.1.2 resulted in 58 identified biomarkers from the Archer dataset, 634 biomarkers from the Brain dataset, and 1040 biomarkers from the Möller-Levet dataset. For each dataset, we fit linear regression models relating the estimated population-level amplitude to population-level measures of MESOR variability, amplitude variability, phase-shift concentration, and residual variance. Across all fitted models and datasets, the coefficient of determination (*R*^2^) exceeded 0.8.

**Figure 4:**
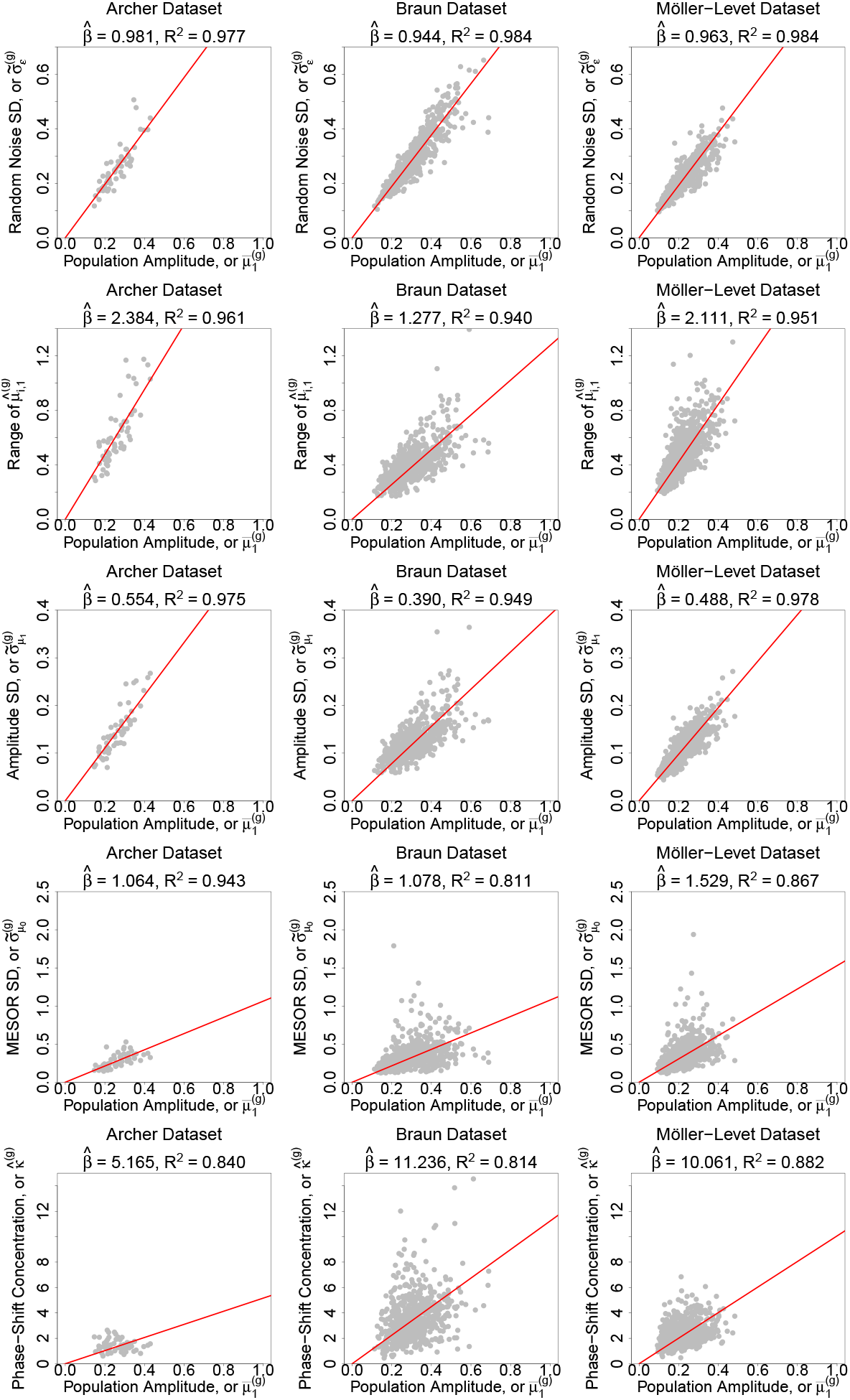
Relationship between average oscillation amplitude and individual-level differences in single-component cosinor model parameters across biomarkers. Points denote biomarkers, lines show least squares fits.

These empirical relationships directly informed the choice of simulation configuration parameters. Regressing the standard deviation of individual-level amplitudes 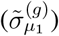 on the corresponding population-level amplitude estimate 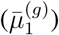 produced values of 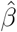 ranging from 0.390 to 0.554 across datasets. Based on this range, we set the standard deviation of the truncated normal distribution used to generate individual-level amplitudes to 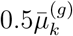 for each harmonic. Similarly, regressing the range of individual-level amplitude estimates on the population-level amplitude estimate produced values of 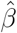 between 1.277 and 2.384. As a consequence, we set the lower and upper bounds of the truncated normal distribution for 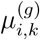 to 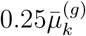and 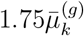, respectively.

For MESOR variability, regressing the standard deviation of individual-level MESOR estimates 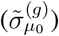 on the population-level amplitude estimates produced values of 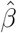 between 1.064 and 1.529. Due to the values of 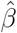obtained, we set the standard deviation of the normal distribution used to generate individual-level MESORs to 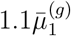.

Finally, regressing the von Mises concentration parameter for individual-level phase-shifts 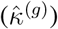 on population-level amplitude estimates produced values of 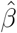 between 5.165 to 11.236. Based on these results, we set the concentration parameter of the von Mises distribution used to generate individual-level phase-shifts to 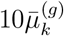.

We note that, in addition to informing the simulation configuration parameter specifications, Figure 4 also shows that regressing the standard deviation of the random noise across individuals 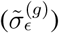 on the population-level amplitude estimates produced values of 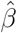 between 0.944 and 0.981. Due to this, we set the standard deviation of the normal distribution used to generate the random noise to 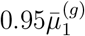.

Figure 5 displays histograms of the population-level amplitude and phase-shift estimates for each dataset. When fitting gamma distributions to the population-level amplitude estimates, the estimated shape parameters ranged from 10.818 to 15.444 and the rate parameters ranged from 34.905 to 59.233 across datasets. As a consequence, we generated population-level amplitudes from a gamma distribution with shape parameter of 12.5 and a rate parameter 45*k* for the *k*th harmonic. The scaling of the rate parameter by *k* implies that the generated amplitudes will typically be smaller for higher-order harmonics, which is consistent with the finding that few genes exhibit statistically significant higher-order amplitudes (Hughes et al., 2009).

**Figure 5:**
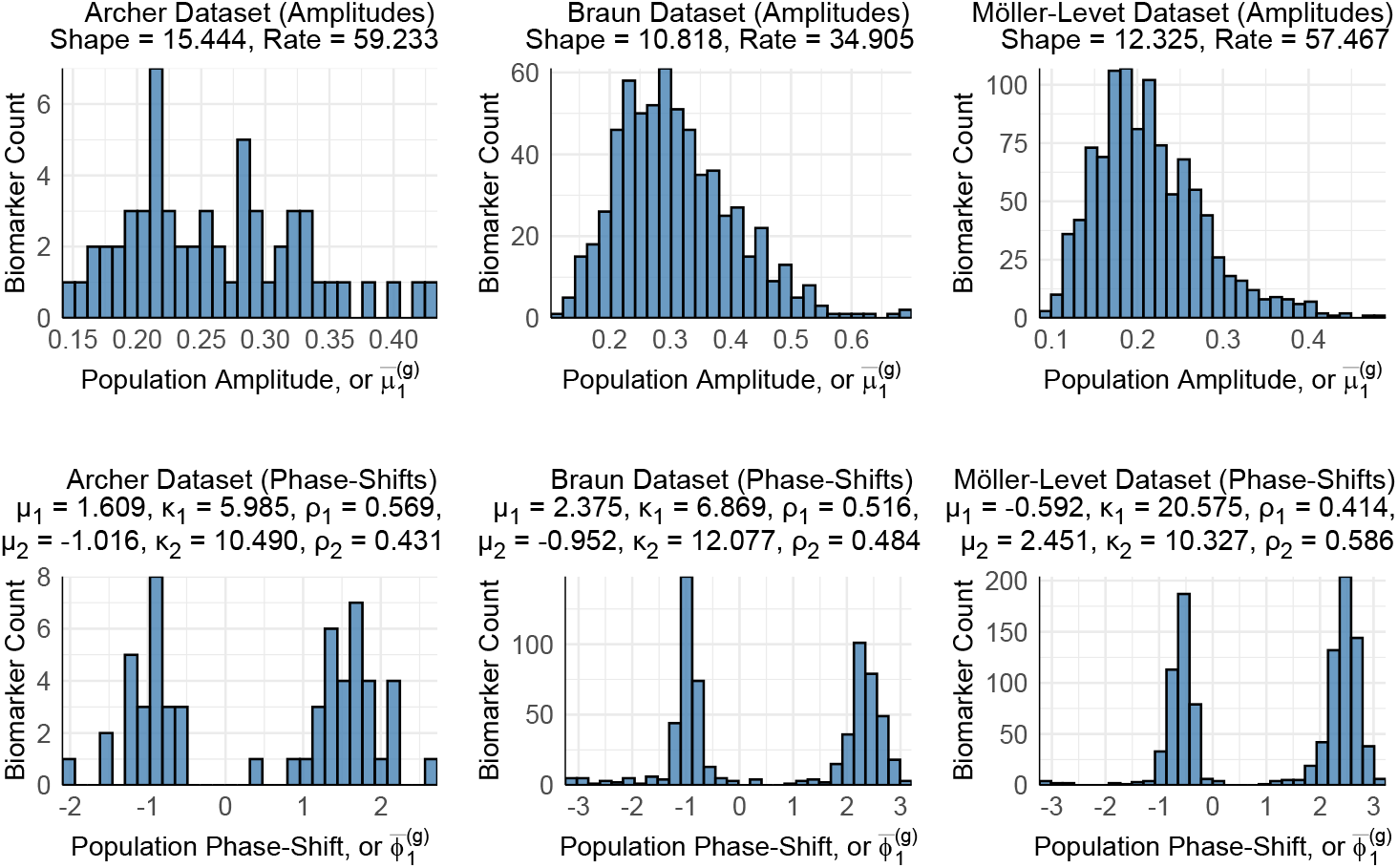
Histograms of estimates for population-level amplitude and phase-shift for the single-component cosinor model with each data set. For the phase-shift histograms, *µ*_*k*_ denotes the mean estimate for the *k*th component, *κ*_*k*_ the concentration estimate for the *k*th component, and *ρ*_*k*_ the mixing proportion for the *k*th component.

For the population-level phase-shifts, we found that the first component of the mixtures of von Mises distributions had mean estimates ranging from -1.016 to -0.592, concentration parameters from 10.327 to 20.575, and mixing proportions between 0.414 and 0.484. The second component had mean estimates ranging from 1.609 to 2.451, concentration parameters from 5.985 to 10.327, and mixing proportions between 0.516 and 0.586. As a consequence, we specified a two-component mixture with equal mixing probability, where there is a 50% chance that a population-level phase-shift is generated from a von Mises distribution with a mean of − *π/*2 and concentration *κ*^(*g*)^ = 7; and a 50% chance that it was generated from a von Mises distribution with a mean of *− π/*2 and a concentration *κ*^(*g*)^ = 14.

## B Correlation Computation

ICT predictions were first aligned to the true ICT so that each prediction was placed within 12 hours of its corresponding true value. Specifically, for each ICT prediction 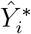 and true ICT 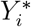 the aligned ICT prediction 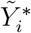 is defined as

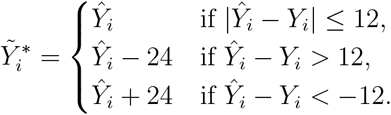

The absolute value of the Pearson correlation between 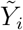 and *Y*_*i*_ was then computed.

